# Whole Plant Transpiration Responses of Common Bean (*Phaseolus vulgaris* L.) to Drying Soil: Water Channels and Transcription Factors

**DOI:** 10.1101/2022.09.23.509270

**Authors:** H. Cordoba-Novoa, B. Zhang, Y. Guo, M.M. Aslam, F.B. Fristchi, V. Hoyos-Villegas

**Affiliations:** Michigan State University, Department of Plant, Insect and Microbial Sciences, 1066 Bogue St, East Lansing, MI, USA; McGill University, Department of Plant Sciences, Montreal, Canada; School of Life Science, Shanxi University, Taiyuan, Shanxi, 030006 China; University of Missouri-Columbia, Division of Plant Science & Technology, 1-31 Agriculture Building, Columbia, MO 65201, USA

**Keywords:** Breeding, abiotic stress, drought, GWAS, transcription factor, aquaporin

## Abstract

Common bean (*Phaseolus vulgaris* L.) is the main legume crop for direct human consumption worldwide. Among abiotic factors affecting common bean, drought is the most limiting. This study aimed at characterizing genetic variability and architecture of transpiration, stomatal regulation and whole plant water use within the Mesoamerican germplasm. A critical fraction of transpirable soil water (FTSWc) was estimated as the inflection point at which NTR starts decreasing linearly. Genome-wide association (GWA) analyses for mean NTR and FTSWc were performed. High variation on mean NTR and FTSWc was found among genotypes. Genomic signals controlling the variation of these traits were identified on *Pv*01 and *Pv*07 some located in intergenic, intronic and exonic regions. A set of novel candidate genes and putative regulatory elements located in these QTL were identified. Some of the genes have been previously reported to be involved in abiotic tolerance in model species, including some of the five transcription factors (TF) identified. Four candidate genes, one with potential water transportation activity and three TFs were validated. The gene *Phvul.001G108800,* an aquaporin SIP2-1 related gene, showed water channel activity through oocyte water assays. Mutant *Arabidopsis thaliana* (*Ath*) lines for the homologous genes of common bean were evaluated in transpiration experiments. Two of the three evaluated TFs, UPBEAT1 and C2H2-type ZN finger protein, were involved in the control of transpiration responses to drying soil. Our results provide evidence of novel genes to accelerate the drought tolerance improvement in the crop and study the physiological basis of drought response in plants.

## Introduction

Pulses are a rich source of dietary fiber, micronutrients, vitamins, carbohydrates, and mono- and poly-unsaturated fatty acids (McCrory et al., 2010). Among pulses, common bean (*Phaseolus vulgaris* L.) is the most cultivated pulse crop for direct human consumption (FAOSTAT, 2019). This crop represents a valuable commodity to ensure food security worldwide, mainly in developing countries. In climate change scenarios, increasing temperatures and the reduction of water availability pose serious threats to food production (Fereres and Soriano, 2007; OECD, 2014). Common bean yield is affected by drought (Beebe et al., 2014), therefore the development of drought-tolerant cultivars is an urgent need in breeding programs (Miklas et al., 2006).

One of the main plant responses to water limitations is reduced transpiration to avoid excessive water loss (Osakabe et al., 2014). Transpiration is limited by partial stomatal closure in response to both increased atmospheric vapor pressure deficit (VPD) and reduction of the fraction of transpirable soil water (FTSW) (Sanchez et al., 2021). An early plant response to drying soil will conserve soil water content late in the season, especially during grain filling, preventing wilting and improving plant drought-resistance (Jafarikouhini et al., 2020; Chiango et al., 2022). The selection of genotypes with improved capacity to maintain water status is crucial to developing new drought-tolerant cultivars (Ray and Sinclair, 1997; Egan et al., 2021). Limited transpiration rates have explained the slow wilting genotype observed in response to drought stress in soybeans (Fletcher et al., 2007). The limited-transpiration trait is now included in commercial soybean (*Glycine max* Merr. L.) (Carter et al., 2016) and maize (*Zea mays* L.) (Gaffney et al., 2015) cultivars for increased yield under water limiting conditions.

The identification of differences in traits such as transpiration among diverse genotypes can be challenging as they show high plasticity across environments in multiple plant species (Ritchie, 1980). Since transpiration of plants is directly related to the soil water content (Kramer, 1944), Ritche (1981) suggested that physiological responses of plants to drying soil should be expressed based on the fraction of extractable soil water content. A methodology to identify patterns in water use is to analyze the normalized transpiration rate (NTR) based on soil water content, and the critical fraction of transpirable soil water (FTSWc). Comparisons based on soil water content are more reliable to detect differences between genotypes attributable to actual variation in plant physiology (Ray and Sinclair, 1997). The FTSWc indicates the plant- transpirable soil water content at which the plant starts closing stomata and reducing transpiration rates. Therefore, FTSWc allows the characterization of genotypes based on their response. FTSW is centered between 0 and 1, where relatively low values of FTSWc indicate that the plant maintains transpiration rates longer and may continue growing under long periods of stress. In contrast, germplasm with relatively high FTSWc thresholds are more sensitive to water deficit and close stomata earlier, making them suitable for intermittent drought stress periods (Egan et al., 2021). High FTSWc may also be evidence for germplasm with faster recovery rates between intermittent drought periods.

The analysis of NTR and FTSWc has been used to characterize stomatal regulation dynamics in maize (Ray and Sinclair, 1997; Devi and Reddy, 2020b); genotypic variation in cotton (Devi and Reddy, 2020a), sorghum (Choudhary and Sinclair, 2014), white clover (Egan et al., 2021); and pearl miller cultivars (*Pennisetum glaucum* (L.) R. Br.) (Kholová et al., 2010a). Despite the challenges of translating the results from greenhouse experiments to the field, the analyses of FTSW and transpiration rate has allowed the characterization of promising genotypes in running breeding programs, being a useful strategy for the study of the plant physiology and advancement of genetic improvement (Fletcher et al., 2007, Gaffney et al., 2015, Chiango et al., 2022). To date, no Genome-wide Association (GWA) studies have been performed to attempt to identify candidate genes involved in the regulation of these constitutive water-conserving mechanisms.

The study of plant transpiration responses to water limitations, along with the identification and validation of candidate genes, provides valuable resources for designing and implementing molecular breeding strategies, such as marker-assisted selection (MAS) for rapid genotype screening, as well as applications in genome editing. The overall objective of this study was to characterize the response of transpiration in Mesoamerican bean accessions to drying soil, identify candidate genes using GWA, and validate key genes involved in the control of transpiration responses.

## Materials and methods

### *Phaseolus* plant material and experiment design

A subset of 82 Mesoamerican common bean genotypes (hereafter drought-tolerant diversity panel, DTDP) from the larger Mesoamerican diversity panel (MDP, 498 individuals) was used in the present study. The DTDP materials were selected for their response to drought in the field; further details on the DTDP are reported in Hoyos-Villegas et al. (2017). The MDP is a diverse set of genotypes from the Mesoamerican gene pool from common bean assembled for genetic studies (Moghaddam et al., 2016). The DTDP was previously selected based on the knowledge from the breeding histories of the genotypes and their performance under rainfed and water limiting conditions within the Bean Breeding Program at Michigan State University (MSU). DTDP is a selected group of varieties, lines, and landraces that differ in their response to drought. More details on the DTDP can be found in Mukeshimana et al. (2014) and Hoyos-Villegas et al. (2017). Three independent greenhouse experiments were conducted at MSU in 2013 (repetitions one and two) and 2014 (repetition three), with a duration 26, 37, and 32 days after treatment (DAT), respectively.

For each repetition trial, treatments consisted of irrigation (control), and drought with three replicates in a randomized complete design. Prior to the beginning of the treatments, pots were saturated and allowed to drain overnight. The next morning, pots were sealed at the bottom and top to avoid any water loss from runoff and evaporation. At the beginning of the experiment the saturated pot weights were determined, and daily pot weights were recorded. Plants under well-watered conditions were replenished with the exact amount of water that had been transpired in a 24-hr period. Drought treatments underwent a progressive dry down with no water replenishment. The drought treatment finished once all the cultivars had died. Further details on this methodology can be found in Ray and Sinclair (1997), Egan et al. (2021), and Chiango et al. (2022).

### Transpiration rate and FTSW

Data from the three experiments were analyzed together (replicates). Transpiration was calculated following the methodology proposed by Ray and Sinclair (1997) by measuring the difference in pot weights every 24 hours. In days when there was no deficit (e.g. due to cloudy conditions between days), values were eliminated for that replicate. The transpiration rate (TR) was calculated as the ratio between the weight of droughted pots and well-watered pots on each day. A second normalization was applied to TR using the mean TR between days two and five (well-watered period) according to the formula:

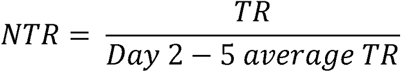

During days 2 – 5, all the plants were still well-watered, as sufficient soil water was available. With this normalization, plants at full transpiration would have a normalized TR (NTR) near to 1.

To determine the extent of water extracted by plants, the fraction of transpirable soil water (FTSW) was calculated as:

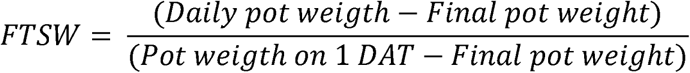

The relationship between NTR and FTSW was obtained by fitting the equation previously described by Muchow and Sinclair, (1991):

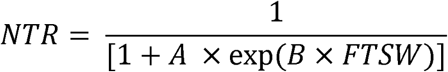

Where A and B are empirical coefficients generated for each genotype through curve fitting. A linear-plateau regression was used to determine the critical FTSW (FTSWc) where NTR starts decreasing linearly and indicates the point at which stomata start to close in stressed plants. FTSWc was determined for each genotype in each replicate. The regressions were done using GraphPad Prism Version 8.0 (GraphPad Software, Inc., San Diego, CA 92121, USA)

## Statistical analysis

The data from the three experiments in common bean were combined and analyzed together in a randomized complete block design (RCBD) with average values across replications within repetitions as replicates. One-way analysis of variance (ANOVA) was conducted using the following model:

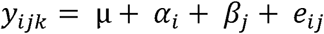

Where µ is the grand mean, *α_i_* is the effect of the *ith* genotype, *β_j_* is the effect of the *jth* repetition experiment, and ### is the random error. Best linear unbiased predictors (BLUPs) were estimated with the restricted maximum likelihood (REML) method using the *lme4* package in R v4.1 (Bates et al., 2015) and BLUP-adjusted means were calculated. BLUP-adjusted means were compared by the least-squares method (α=0.05). Broad-sense heritability (H^2^) was estimated by:

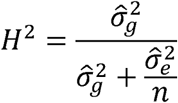

Where 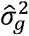 is the genotypic variance, 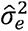 is the error variance, and n is the number of replicates (experiments).

## Genome-Wide Association Analyses

### Genotypic data

The DTDP was previously genotyped (Moghaddam et al., 2016). The genotypic data contains around 200K single nucleotide polymorphisms (SNPs) and can be found on the BeanCAP website (http://arsftfbean.uprm.edu/beancap/research/, Accessed Jun. 2022). SNPs with a minor allele frequency (MAF) < 0.02 were filtered out. This threshold was chosen to facilitate the identification of rare genotypes. Linkage disequilibrium (LD) between markers was calculated using all the SNPs within a sliding window of 50 SNPs in TASSEL 5 software (Bradbury et al., 2007).

The LD decay pattern for each chromosome and whole genome was analyzed by nonlinear regression fitting the expectation of the parameter r^2^ and the distance between each pair of markers (Hill and Weir, 1988; Remington et al., 2001). Population structure was evaluated using the variational Bayesian inference in fastSTRUCTURE (Raj et al., 2014) with two populations (K = 2) based on previous knowledge of the subpanel (Hoyos-Villegas et al., 2017).

### Genomewide Association Analysis (GWAS)

A GWAS analysis was performed using 182,243 SNPs with a MAF > 0.02, and the BLUP-adjusted means of the FTSWc and mean NTR under water deficit for each genotype. Analyses were carried out using multiple models, including Mixed Linear Model (MLM), compressed MLM (CMLM), Settlement of MLM Under Progressively Exclusive Relationship (SUPER), Fixed and random model Circulating Probability Unification (FarmCPU), and Bayesian-information and LD Iteratively Nested Keyway (BLINK) implemented in GAPIT v.2 (Tang et al., 2016). Individual relatedness was also considered with the kinship matrix that is automatically calculated within GAPIT using the VanRaden method (VanRaden, 2008). Peak SNPs with *p-*values above a Benjamini-Hochberg false discovery rate (FDR) threshold of 0.05 were considered significant and were further examined. The variance explained by significant SNPs was estimated by the coefficient of determination (*R^2^*) of a linear regression model. The model included all the five PCs from PC analysis (PCA), the SNP marker parameterized in accordance with the gene action determined using the general, additive, and simplex dominance models in the R package GWASpoly (Rosyara et al., 2016), and a vector of random residuals (Wallace et al., 2016).

## Haplotype analysis and candidate genes

Haplotype blocks were defined using the solid spine of LD method implemented in Haploview v.4.2 software with default parameters (Barrett et al., 2005). The proportion of the phenotypic variance explained by significant SNPs from GWAS (*R^2^*) was recalculated after including the SNPs present in the same haplotype block.

To identify candidate genes, JBrowse on Phytozome v13 (Goodstein et al., 2012) was used to browse the common bean reference genome v.2.1 (Schmutz et al., 2014) within a ± 100- Kb window centered on each associated SNP. The search window for candidate genes was defined based on the lower end of the calculated LD decay and to be consistent with previous studies in common bean (M oghaddam et al., 2016; Jain et al., 2019; Raggi et al., 2019). Candidate genes were selected based on their annotation and literature review related to drought and other abiotic stresses.

## Validation of candidate genes

Based on the gene model annotation and the potential role in the water status of the plant, four candidate genes were selected for further validation. A gene with potential water channel activity was validated using oocyte swelling assays. For other genes, due to the challenges associated with common bean transformation and regeneration such as recalcitrance and lack of repeatability (Yilmaz et al., 2022), *Ath* mutant lines with knockouts in the homologous genes to those identified in common bean were used.

### In vitro RNA synthesis and oocyte swelling assays

The gene DNA sequence with Gateway attachment sites was *in-vitro* synthesized (Sangon Biotech [Shanghai] Co., Ltd.). The gene was inserted into the pDONR207-Dest vector by BP reactions using BP Clonase II (Invitrogen) and verified via sequencing (Sangon Biotech, Shanghai, China). For *Xenopus laevis* oocytes water transport assays, the gene was inserted into the pGT-nHA-Dest vector (Zhang et al., 2015).

For the oocytes water transport assays, plasmids were linearized, and the capped cRNA was synthesized *in-vitro* using the mMESSAGE mMACHINE T7 High Yield Capped RNA Transcription Kit (Ambion, USA) as previously reported (Zhang et al., 2015). The cRNA quality was checked with agarose gel and quantified in a Nano-300 spectrophotometer (Hangzhou Allsheng Instruments, China).

Oocyte water transportation assays were conducted as previously reported (Canessa Fortuna et al., 2019). Stage VI oocytes were isolated from mature *X. laevis*, and the follicular cell layer was digested with 2 mg/mL collagenase (Sigma-Aldrich) for 20 min before injection. Injected oocytes were incubated in ND96 (96 mM NaCl, 2 mM KCl, 1 mM MgCl2, 1 mM CaCl2, and 10 mM HEPES-NaOH, pH 7.4) supplemented with gentamycin (5 mg/L) at 18°C for three days before recordings. The oocytes’ osmotic water permeability coefficient (P_f_) was determined by measuring the oocyte swelling rate in response to ND96 buffer diluted four-fold with distilled water. The changes in oocyte volume were video-monitored by an SZ680 zoom stereo-microscope with a color camera (Chongqing Optec Instrument, China). Cell swelling was video-captured in still images, and the oocyte was treated as a growing sphere whose volume could be inferred from its cross-sectional area (Image J software version 1.37, NIH, USA). P_f_ was calculated according to previous reports (Canessa Fortuna et al., 2019; Fox et al., 2020; Baena et al. 2024). P_f_ = V_o_[d(V/V_o_)/d*t*]/[S V_w_ (Osm_in_ – Osmo_ut_)], where V_o_ is the oocyte volume (9 x10^-4^ cm^3^), V/V_o_ is the relative volume, S is the surface area of the oocyte (0.045 cm^2^), V_w_ is the molecular volume of water (18 cm^3^ mol^-1^), and Osm_in_-Osm_out_ is the osmotic driving force. Non-injected oocytes were used as negative controls. All osmolarities were measured in the vapor pressure osmometer (Vapro 5600 Wescor, USA).

The presence of the proteins in oocytes was confirmed by western blot. Samples were separated by SDS-PAGE on 10% gels. Immunoblot analysis was performed after transfer to nitrocellulose filters and blocking using both primary commercial antibodies α-HA (1:10000, Abcam) and the secondary goat anti-rabbit antibodies (Abcam). ECL detection reagent (Yeasen, Shanghai, China) was used for the detection.

### Ath mutant lines transpiration experiments

Mutant lines with knockouts for the homologous of the candidate genes were obtained from the Arabidopsis Biological Resource Center (ABRC) of The Ohio State University. Seeds were increased and confirmed by PCR with primers designed following the protocol for SALK T-DNA (Table S1). *Ath* transpiration experiments followed the same procedure outlined in the above section, with minor modifications to *Ath* as per Wahbi et al. (2007). Briefly, seeds were sterilized using 70% ethanol and then rinsed twice with sterile distilled water. Seeds were placed in Petri dished with moist filter paper, wrapped in aluminum foil and incubated for three days at 4°C to break dormancy. Seeds were sown in single pots with commercial substrate (Agro mix® G7) and covered for two days with plastic wrap. Once the seeds emerged, they were carefully transplanted into single pots with a volume of about 73.4 cm^2^ and the same G7 substrate.

Two independent experiments were run under growth chamber conditions with artificial light of 230 – 250 mol m^−2^ s^−1^, a photoperiod of 12 h, an average temperature of 22°C and a humidity of 60%. Each experiment consisted of five replicates per line and two treatments: Drought and irrigated. The genotype Columbia was used as the wild type (WT). Treatments started 10 days after transplanting. Pots were overwatered and allowed to drain overnight. The following morning, the top and bottom were carefully sealed with plastic wrap and parafilm to avoid water loss, and the initial pot weight was recorded. For plants under drought conditions, irrigation was suspended, and daily weights were recorded. Control plants (irrigated) were maintained at 75% of the pot capacity. Data was analyzed following the same methodology as in common bean experiments with an RCBD mentioned before. To analyze the differences among mutant lines, the slope of the linear fraction of the regression and the mean of the NTR after the FTSWc values were obtained.

## Results

### Transpiration rate response to drying soil and heritability

The average NTR per genotype during the evaluation period ranged from 0.33 in Yolano to 0.96 in ABCP-8, with a global mean of 0.52 (Table 1), a coefficient of variation (CV) of 0.26 and a H^2^ of 0.53. There was a consistent relationship between NTR and FTSW. The NTR value decreased with a linear tendency below the FTSWc for each genotype. The FTSW threshold (FTSWc) varied from 0.23 in the pink bean genotype Harold to 0.99 in the black bean T-39 with an overall mean of 0.45, a CV of 0.47, and significant differences between the maximum and the minimum (*p*<0.05; Table 1). Figure 1 depicts the response of NTR to a decreasing FTSW for the bottom (mean FTSWc = 0.26; Figure 1A) and top five (mean FTSWc = 0.71; Figure 1B) genotypes. In the linear regression fraction post FTSWc for the group of genotypes, the slope was lower (1.36 ± 0.13) in those genotypes with a higher FTSWc compared to the slope (3.49 ± 0.47) for the genotypes with a low FTSWc value (Figure 1). Using the genotypic and error variances, the estimated H^2^ for FTSWc was 0.38.

**Figure 1.**
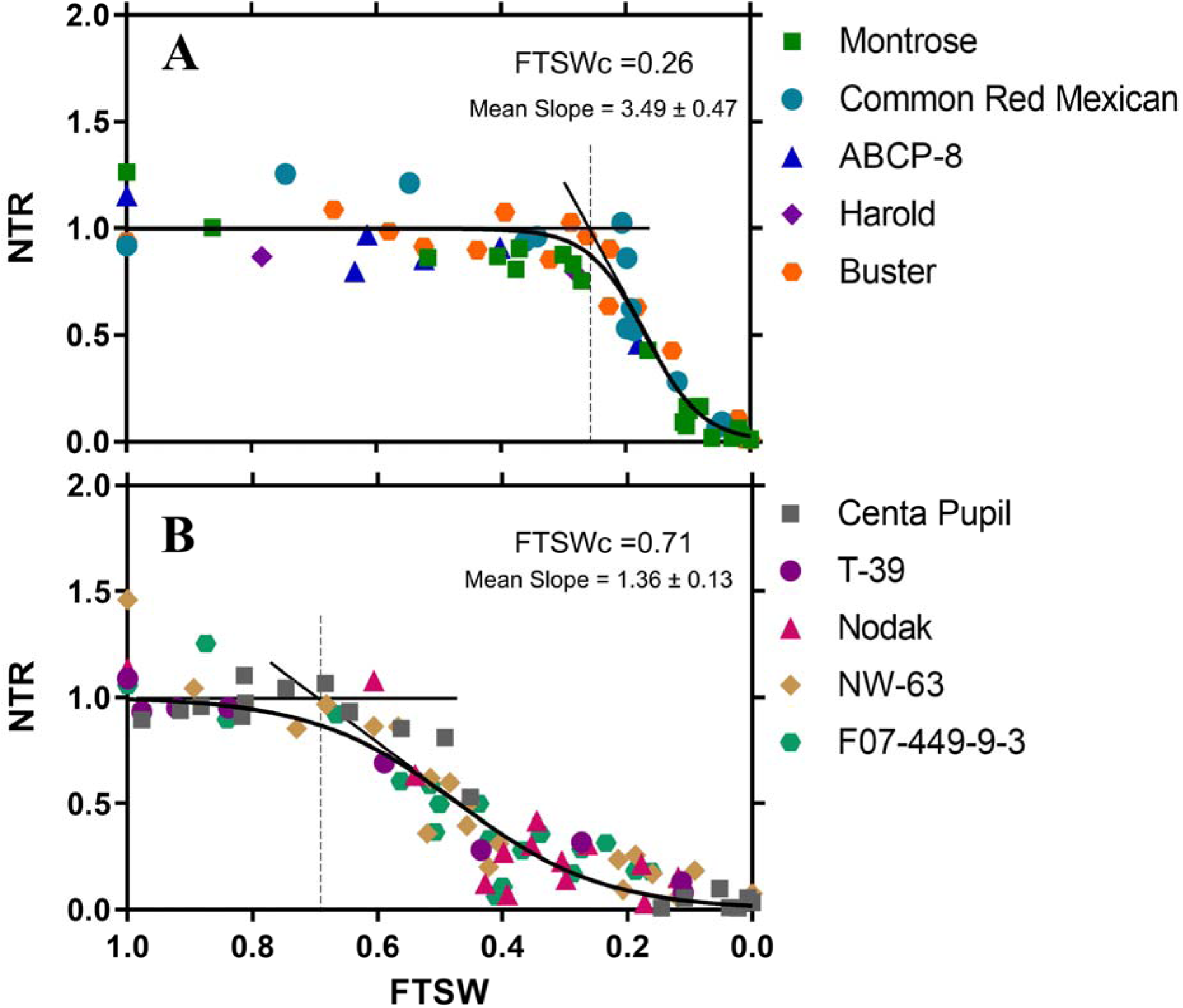
Mean normalized transpiration rate (NTR) in response to the fraction of transpirable soil water (FTSW) in the five genotypes with the lowest FTSWc (A) and the five genotypes with the highest FTSWc (B). Mean values for FTSWc and the slope (mean Slope ± SE) of the linear regression post-FTSWc are shown.

**Table 1.**
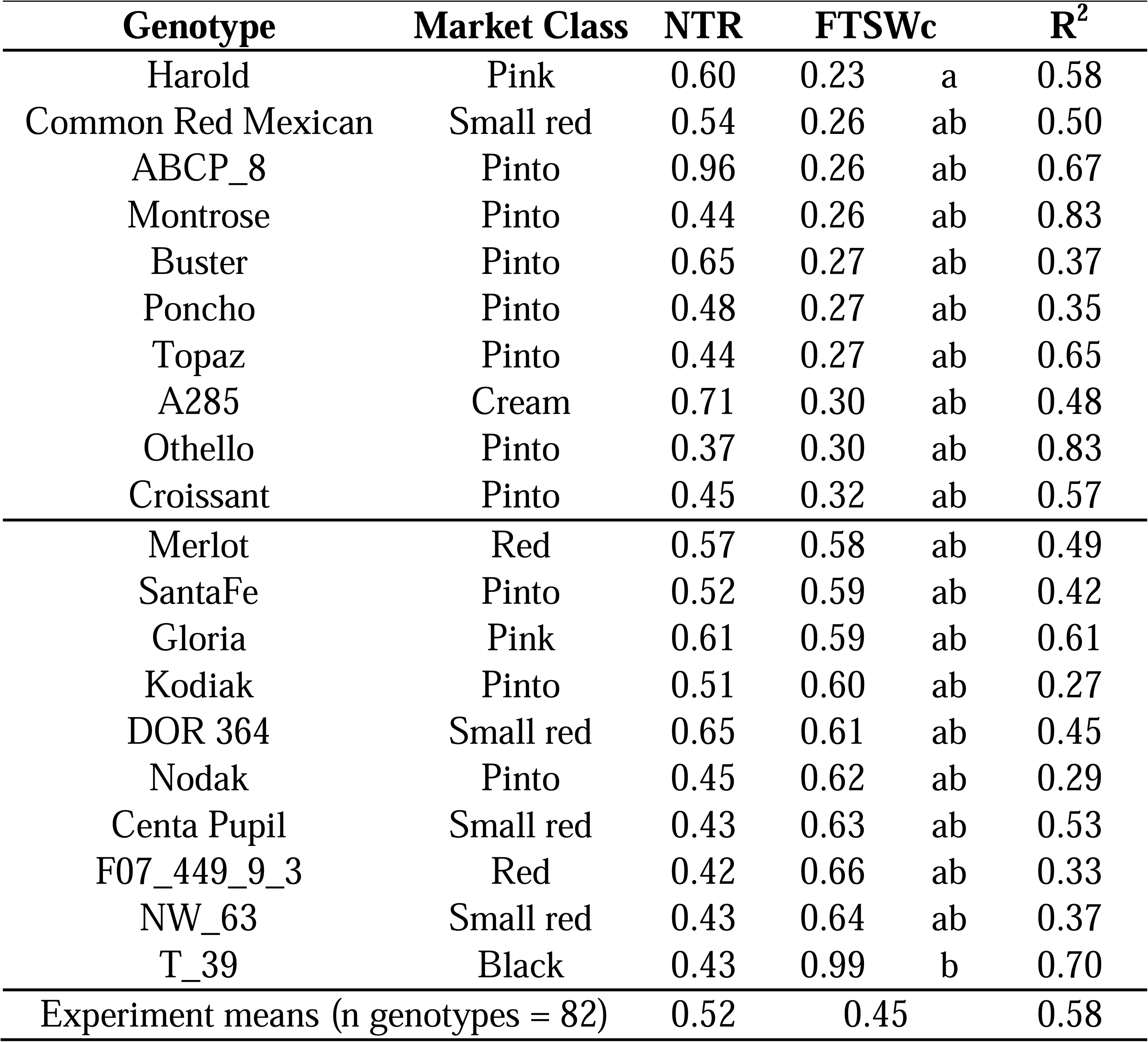
Normalized Transpiration Rate (NTR) and critical Fraction of Transpirable Soil Water (FTSWc) for the top and bottom ten evaluated common bean genotypes. The R^2^ indicates the linear-plateau regression fit to the data.

### Population structure and LD decay

Two subpopulations were identified in the DTDP (K=2) with varying probabilities of ancestry among genotypes. Admixture between the two subpopulations was also observed. The chromosome-wise analysis of the LD decay pattern showed that in the DTDP, LD decayed at a range from 135.5 Kbp in *Pv04* to 848.7 Kbp in *Pv09* with a genome-wise decay distance of 363.3 Kbp.

### Genome-Wide Association and haplotype analysis

Based on the adjustment of the models in the QQ plots, all the tested models performed similarly with a slight improvement in Blink for FTSWc (Figure S1) and FarmCPU for mean NTR (Figure S2). Blink and FarmCPU were used for the analysis of marker-trait associations for FTSWc and mean NTR, respectively. FarmCPU and Blink have also shown improved statistical power with both simulated and real datasets (Huang et al., 2019; Kaler et al., 2020).

In FTSWc, one SNP was identified on *Pv01* at 33,516,386 bp (C/T; *Pv01*FTSWc) and another on *Pv07* at 9,224,979 bp (C/T; *Pv07*FTSWc) (Figure 2A). Each SNP for FTSWc had an effect of +0.26 units and explained the 31% of the phenotypic variation. For mean NTR, two significant SNPs were identified in *Pv01* at 28,451,384 bp (C/A; *Pv01*NTR_1) and 35,489,136 bp (A/G; *Pv01*NTR_2) with an effect of -0.22 and +0.19 units, respectively (Figure 2B). *Pv01*NTR_1 explained the 23% of the phenotypic variation (R^2^), and *Pv01*NTR_2 the 24%. The four SNPs were detected using the multiple models and had a MAF of 0.02.

**Figure 2.**
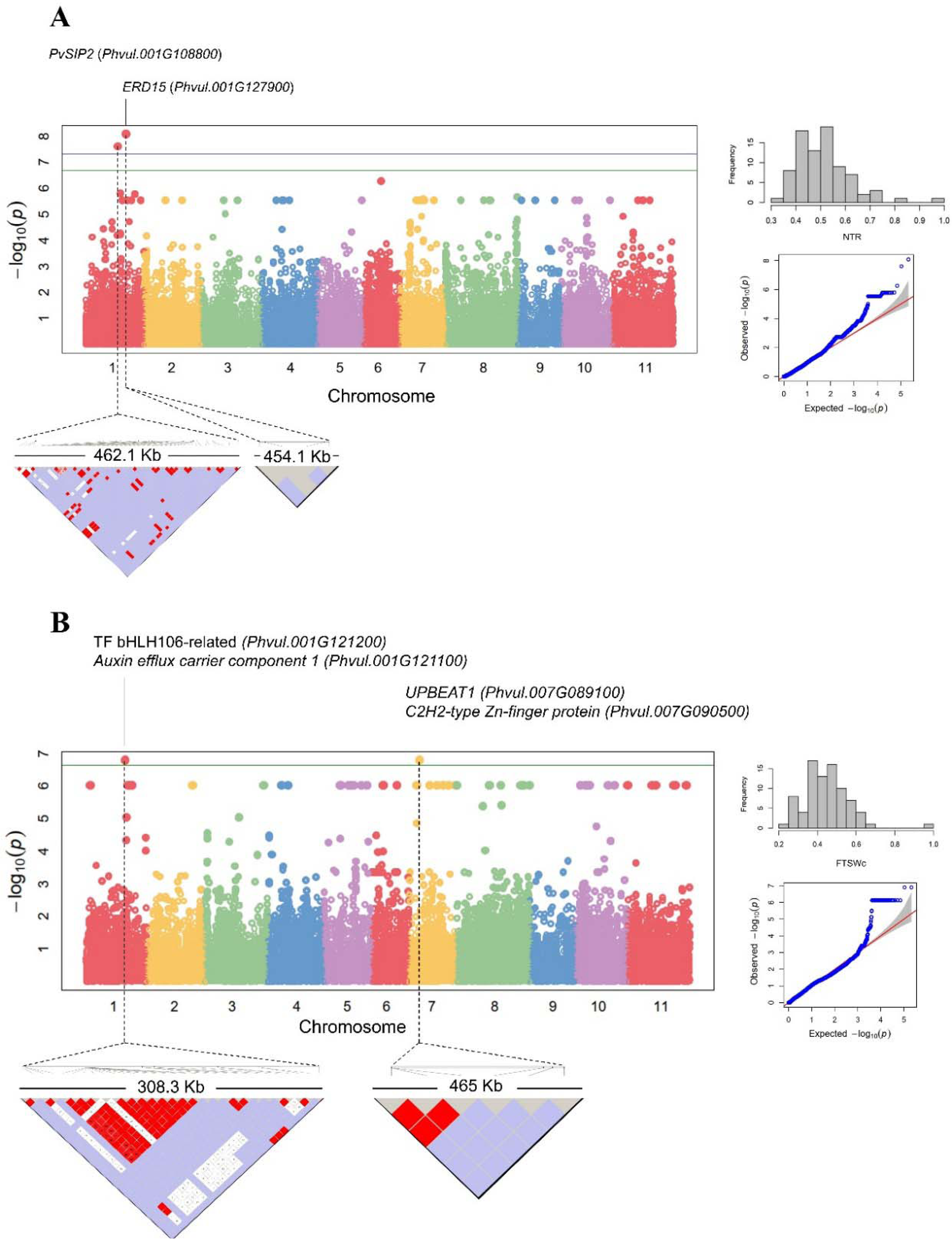
Manhattan plots of QTL identified for NTR (A) using FarmCPU model, and for FTSWc (B) using Blink model in the subpanel of 82 Mesoamerican beans. SNPs above the FDR- corrected threshold were considered as significant. The haplotype block for each significant SNP is depicted in the bottom of each Manhattan plot. The figures to the right of the Manhattan plots are the phenotype distribution histogram and the Q-Q plot with the adjustment of the model for each trait.

The genotype T-39 that had the highest FTSWc was homozygous for the alternative allele (TT) of *Pv01*FTSWc and *Pv07*FTSWc. NW-63 was heterozygous for the significant SNPs of FTSWc and was among the top five values. Harold, lowest FTSWc (0.23) was homozygous for the reference alleles (CC). For *Pv01*NTR_1, ABCP-8 is homozygous (AA) for the alternative allele and had the highest mean NTR (0.96), followed by A-55 which is heterozygous (C/A).

ABCP-8 and A-55 are homozygous (GG) for the alternative allele of *Pv01*NTR_2 and Yolano is homozygous for the reference allele (AA) and had the lowest mean NTR.

In the haplotype analysis for Pv01, the SNP *Pv*01FTSWc was grouped with 29 additional SNPs in a haplotype block of 308 Kbp. Out of the 30 SNPs in the haplotype, 15 were in genic regions and the others in intergenic regions. The haplotype containing *Pv*01FTSWc explained a higher variation (42%) compared to the single SNP (31%). On mean NTR, *Pv01*NTR_1 was grouped with 48 additional SNPs where the marker ss01_28185985 is in a coding region. The block had a size of 462 Kbp and explained a higher total phenotypic variation (36%) compared to the single SNP (23%). *Pv01*NTR_2 was in a block of 454 Kbp with four SNPs, but no increases in the R^2^ were observed for the haplotype compared to the individual significant SNP.

For haplotypes on Pv07, *Pv07*FTSWc was grouped with 6 additional markers in a haplotype of 464 Kbp with the SNP ss07_9279035 located on a genic region. For this haplotype, no increases in the variation explained were detected.

### Candidate genes

The high homozygosity present in common bean due to its predominantly self-pollinating mating system leads to high estimates of LD thresholds in different populations. The estimated LD decay for the DTDP showed wide variation between chromosomes (135.5 Kbp to 848.7 Kbp). To reduce the rate of false positive candidate genes and based on previous studies, including the MDP estimations (Moghaddam et al., 2016; Hoyos-Villegas et al., 2017), we used a 100 Kbp window for candidate gene identification. A total of 23 genes were identified for the four haplotype blocks containing the significant SNPs. Four genes were associated with the haplotype of *Pv*01FTSWc, 12 with the haplotype for *Pv07*FTSWc, three genes with the haplotype containing *Pv*01NTR_1, and four with the haplotype containing *Pv*01NTR_2. However, considering their annotations and potential role in abiotic stress responses, genes were filtered and a total of 14 genes were kept as potential candidates, seven on *Pv01*, and seven on *Pv07* for limited transpiration in common bean (Table 2). Interestingly, four of the genes are annotated as transcription factors (TFs). A TF on *Pv01* was bHLH106-related (*Phvul.001G121200*); and on *Pv07*, TFs were UPBEAT1 (*Phvul.007G089100*), No apical meristem (NAM) protein (*Phvul.007G089600*), and C2H2-type Zn-finger protein (*Phvul.007G090500*).

**Table 2.**
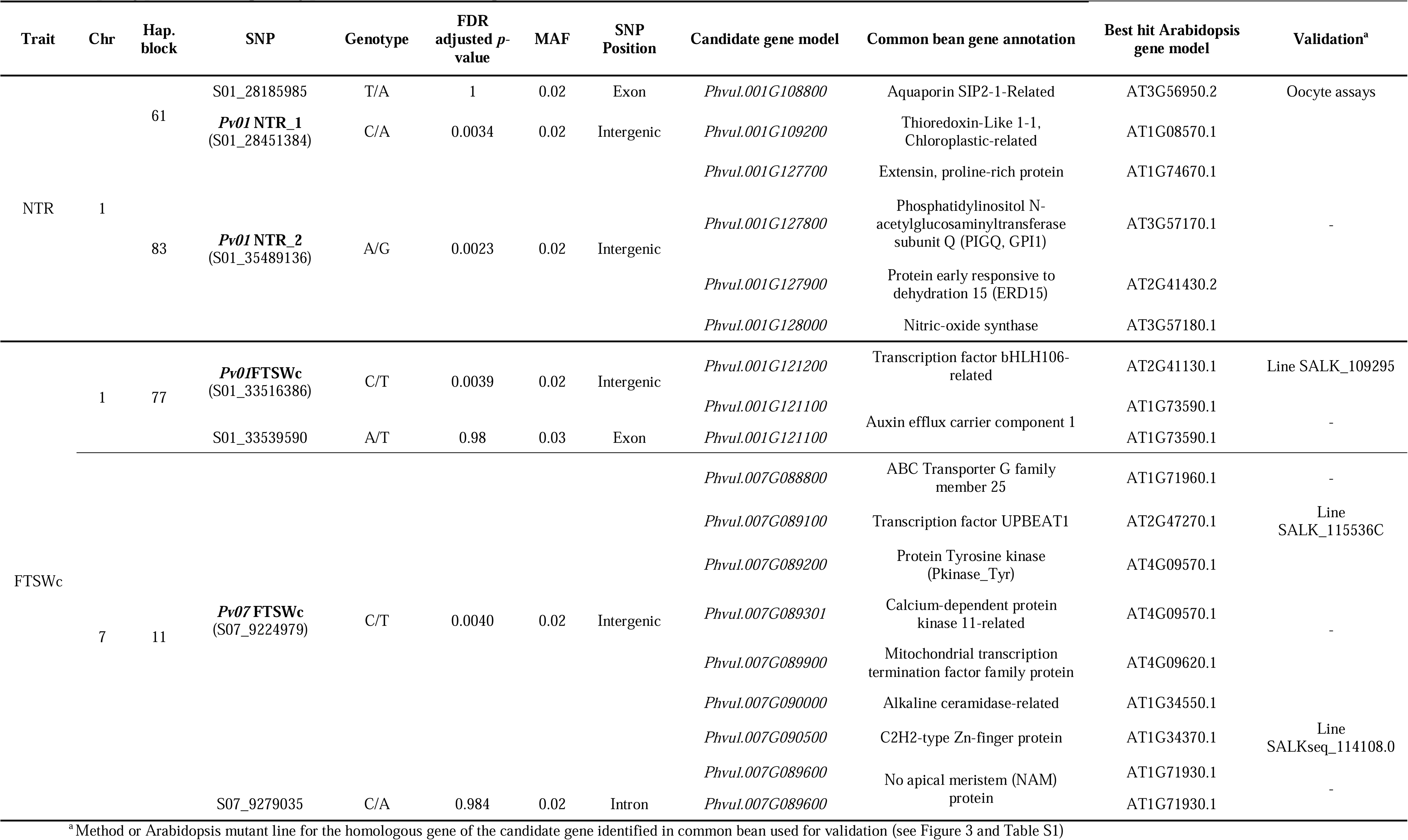
Significant associations for Normalized Transpiration Rate (NTR) and critical Fraction of Transpirable Soil Water (FTSWc). Haplotype blocks, genotypes, and candidate genes for each SNP are indicated.

### Oocyte water transport assays

The candidate gene Phvul.001G108800 encodes an Aquaporin SIP2-1-Related (PvSIP2) protein, which is implicated in potential water transport activities. To demonstrate the functionality of this aquaporin, we carried out the oocyte swelling experiment according to previous literature (Canessa Fortuna et al., 2019; Baena et al., 2024). Our findings revealed that expression of PvSIP2 in oocytes enhanced osmotic water permeability and promoted swelling as compared to control oocytes only injected with water (Figure 3A). Moreover, higher doses of PvSIP2 cRNA injection resulted in a significant increase in osmotic water permeability and greater cellular volume swelling (Figure 3B). The presence of the protein in the oocytes was confirmed, where higher concentrations of the gene corresponded to more pronounced band intensities (Figure 3C).

**Figure 3.**
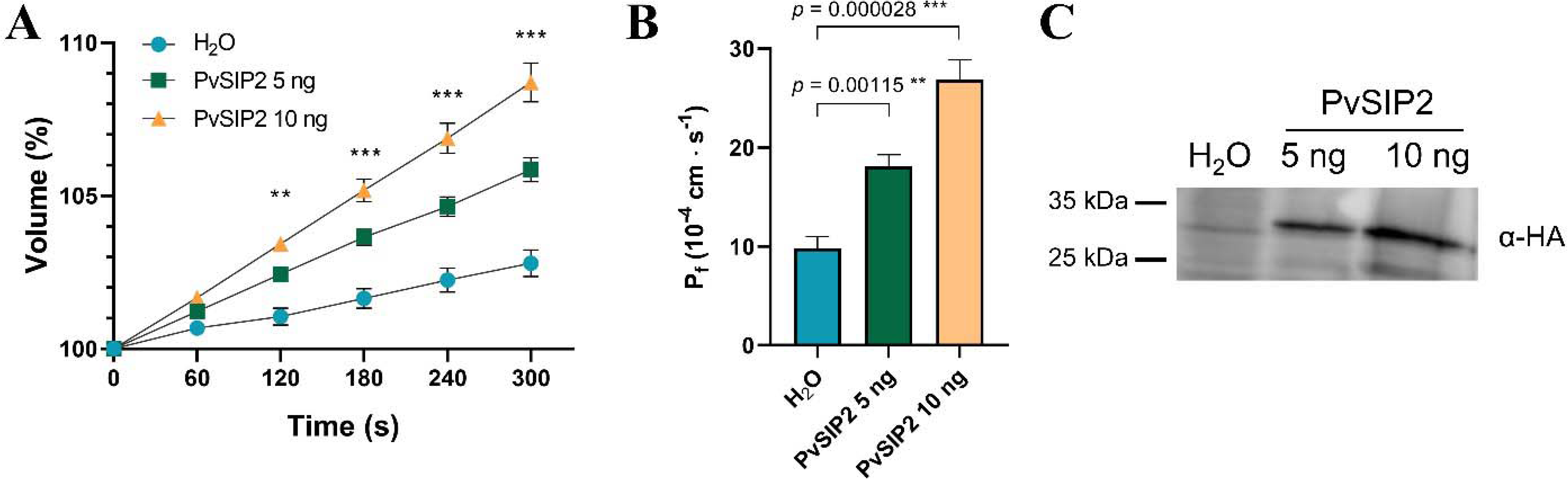
*Phvul.001G108800* (PvSIP-2) has aquaporin channel activity. Time course of the relative volume change in *Xenopus oocytes* injected with cRNA for the expression of PvSIP-2. The oocytes were imaged before injection (0 sec) and after injection with cRNA at 60, 120, 180, 240 and 300 sec (A). Osmotic water permeability coefficient (P_f_) measured on the imaged oocytes (B). Immunoblots detecting the expression of PvSIP-2 protein (approx. 29 kDa) in oocytes (C). The oocytes injected with water served as control. Data are means ± SE, n = 20-26 oocytes expressing PvSIP-2, using cRNA 5 ng/oocyte and 10 ng/oocyte. Statistical significance is indicated by * (p<0.05), ** (p < 0.01) and *** (p < 0.001).

### Ath mutant lines experiments

Three mutant lines SALK_109295 (*AT2G41130*), SALKseq_114108.0 (*AT1G34370*), and SALK_115536C (*AT2G47270*) were evaluated for the homologous genes of *Phvul.001G121200*, *Phvul.007G090500*, and *Phvul.007G089100,* respectively (Table 2). In the lines and the WT, the NTR and FTSW showed the same relationship as observed in the bean experiments (Figure 4 A-D). Despite no significant differences in the FTSWc values, the line *AT2G41130* had a FTSWc (0.37) similar to the WT (0.32). The line *AT1G34370* had a lower FTSWc (0.24) and the line *AT2G47270* had a higher value (0.42) compared to the control.

**Figure 4.**
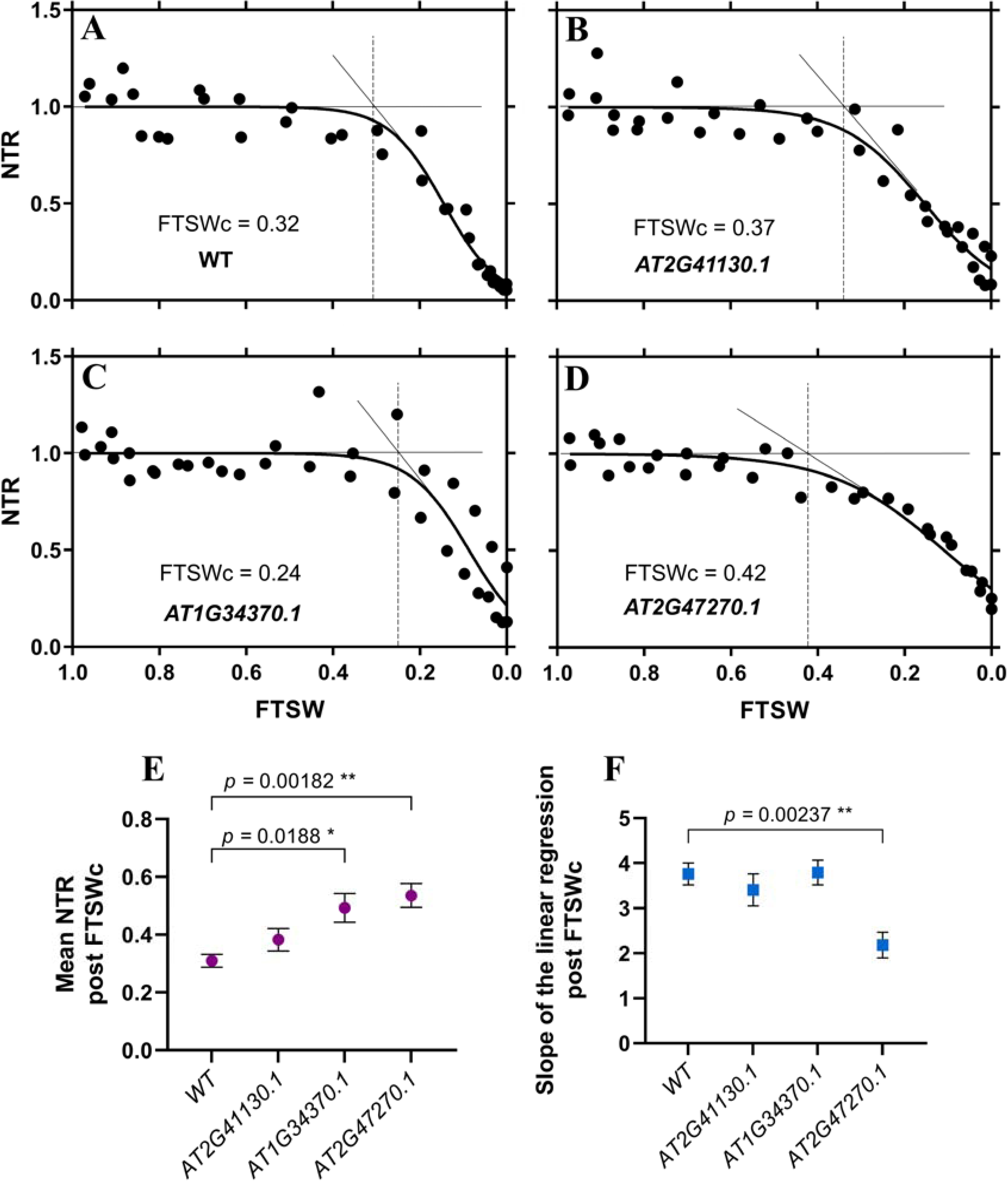
Mean normalized transpiration rate (NTR) in response to the fraction of transpirable soil water (FTSW) in the Columbia WT line (A) and three mutant Arabidopsis lines for the homologous candidate genes identified in common bean (B-D). Line SALK_109295 (*AT2G41130*) for candidate gene *Phvul.001G121200* (B), SALKseq_114108.0 (*AT1G34370*) for candidate gene *Phvul.007G090500* (C), and SALK_115536C (*AT2G47270*) for candidate gene *Phvul.007G089100* (D). Mean NTR post-FTSWc in the WT and evaluated lines (E), and slope of the linear regression post-FTSWc for WT and evaluated lines from A – D (F). Statistical significance is indicated by * (p<0.05), ** (p < 0.01) and *** (p < 0.001) in E and F.

The lines *AT1G34370* and *AT2G47270* had significantly higher mean NTR after the FTSWc (0.49 and 0.53, respectively) with respect to the WT (0.31; Figure 4E). In the slope of the linear fraction of the regression post FTSWc, as a way to measure the rate of the decline of the NTR in response to the drying soil, the line *AT2G47270* had a significantly lower rate (2.18) compared to the WT (3.76; Figure 4F).

## Discussion

The present study investigated the transpiration rate response of common bean and genotypic differences among Mesoamerican accessions. Under drought-like conditions, stomata closure is among the first physiological responses to avoid water loss through transpiration (Osakabe et al., 2014). Stomatal closure under drought is mainly triggered by an increase in the abscisic acid (ABA) concentration in leaves (Bharath et al., 2021). However, it can also be induced by hormones like salicylic acid (SA), methyl jasmonate (MJ) and ethylene; metabolites such as proline and polyamines; and gaseous signals such as hydrogen sulfide (H_2_S) (Agurla et al., 2018). Stomatal closure triggered by these components is mediated by hydrogen peroxide (H2O_2_), calcium (Ca^2+^), and nitric oxide (NO) (Agurla et al., 2018). Likewise, drought causes the ABA-dependent and independent activation of key transcription factors (TFs) (Nakashima et al., 2014).

Stomatal limitations restrict gas exchange rate, photosynthetic activity, and plant growth and development. As a response to drought, tolerant varieties can be identified based on a slow leaf wilting phenotype. The soybean cultivar PI 471938 is an example of a slow wilting genotype that exhibits water-conservative traits by maintaining TR (Fletcher et al., 2007). A reduced or capped TR under water-limited conditions is usually observed in drought-tolerant genotypes compared to susceptible materials in short drought periods (Fletcher et al., 2007; Kholová et al. 2010). Reductions in TR are associated with early stomatal closure and high FTSW thresholds (Ray and Sinclair, 1997; Egan et al., 2021). For the soybean cultivar PI 471938, it has been ruled out that its resistance is due to changes in the FTSWc (Sadok et al., 2012), but the TR does not increase compared to susceptible cultivars under water stress (Fletcher et al., 2007). Implying specific responses for TR and additional mechanisms involved in water conservation that may not be fully detected by sole differences in the critical FTSW.

Among the genotypes with the lowest mean NTR (Table 1), Othello has been reported as a drought-resistant cultivar, but US-1140 and UI-59 as low-yielding cultivars (Muñoz-Perea et al., 2006), which suggests that the stomatal responses of common bean to water deficit have varying impacts on plant performance and is influenced by other traits depending on the cultivar.

A lower TR indicates that the plant favors a conservative water use mechanism and survival for longer periods (Kholová et al. 2010). Devi and Reddy (2020) observed a positive correlation between TR and FTSWc when studying different cotton accessions. In contrast, we observed no relationship between mean NTR and FTSWc since the genotypes with the lower mean NTR did not have a higher FTSWc (Table 1). This can be explained by genotypic differences in stomatal and hydraulic conductance among the evaluated genotypes. Hydraulic conductance affects stomatal conductivity, transpiration rate, and water potential, and its variation depends on anatomic changes in membrane thickness, structure, and permeability (Hacke and Way, 2014; Simonin et al., 2015). Adjustments in hydraulic conductance may also depend on changes in water transport in roots and shoot via aquaporins (Choudhary and Sinclair, 2014). In French bean leaves, Sôber, (1998) observed that high stomatal conductance rates were accompanied by a high hydraulic conductivity. Additionally, Choudhary and Sinclair (2014) found that differences in TR among sorghum genotypes result from a varying hydraulic conductance where some genotypes with low TR can maintain high rates of both stomatal and hydraulic conductance. The slow wilting soybean PI 471938 has been found to have a limited hydraulic conductance in leaves while maintaining constant TR (Sinclair et al., 2008). It is possible then, that the genetic variation in the evaluated panel allows some cultivars to have an early stomatal closure (high FTSWc) with an improved transpiration and hydraulic conductance. An early stomatal closure as a water conservative trait will also result in slower reductions in TR after the FTSWc as observed in the linear regression slope (Figure 1). In contrast, low FTSWc may provide a plant with the ability to conserve water under intermittent drought conditions, but this must be coupled with an equivalent rate of stomatal reopening upon rehydration. In addition, growth and development (as they pertain to seed yield) must not be severely compromised, placing further emphasis on efficiencies between photosynthetic rate and CO_2_ exchange under intermittent and terminal drought conditions.

Plants with high TRs may keep stomata open longer in drying soil, which results in lower FTSWc values (Ray and Sinclair, 1997). However, when the FTSW reaches critical values, the reduction of TR will be more drastric (higher slope). Genotypes with low FTSWc maximize water use and are desirable for short-term water deficit periods (Egan et al., 2021). Harold had the lowest threshold (0.23) and has been reported to have a moderate to high level of resistance to drought (Singh et al., 2001). Common Red Mexican (FTSWc=0.26) is a landrace that exhibits high level of water stress tolerance (Muñoz-Perea et al., 2007), despite having a moderate mean NTR in the present study (0.54). As explained above, a varying dynamic between TR and FTSWc may be explained by genetic differences in stomatal and hydraulic conductance. We did observe that ABCP-8 had the highest mean NTR (0.96) and one of the lowest FTSWc (0.26; Table 1). ABCP-8 is a pinto bean line bred for common bacterial blight resistance by crossing the donor XAN 159 and the recurrent parent ‘Chase’ pinto (Mutlu et al., 2005). It is worth noting that the line XAN 159 was developed using interspecific crosses with tepary bean (*Phaseolus acutifolius* A. Gray) (Thomas and Giles Waines, 1984), which has been referred as a source of resistance for heat and drought stress (Souter et al., 2017; Moghaddam et al., 2021). It is possible that genetic elements involved in the response of the cultivar to drying soil and response to drought were introgressed into ABCP-8 from tepary bean accessions.

Drought tolerance is a complex trait with high variation in multiple structural, physiological, and metabolic components (Bowles et al., 2021). Similar to our results, previous studies on drought responses in common bean have identified QTL on Chr01 and Chr07 (Mukeshimana et al., 2014; Briñez et al., 2017; Dramadri et al., 2019). In a biparental population of Andean genotypes, Sedlar et al. (2020) identified QTL for water potential mainly in linkage groups one and seven. Leitão et al. (2021) found significant associations for stomatal conductance under different water regimes in Portuguese accessions. However, they found no associations on Chr07, and one SNP was located on Chr01 in a different locus compared to our results.

Depending on the breeding purpose, once validated, the identified SNPs can be used for marker-assisted selection (MAS). Different types of markers for associated SNPs such as Kompetitive Allele Specific PCR (KASP) markers can be designed to select parents and offspring carrying the desired superior alleles for drought tolerance (Kelly and Bornowski, 2018). Furthermore, candidate genes and their corresponding homologous in related species are new sources of information for biotechnology approaches such as transgenesis and genome editing.

For short drought periods, selections favoring the alternative alleles (T) for *Pv01*FTSW and *Pv07*FTSW would increase the FTSWc and allow an early stomatal closure. In contrast, if breeding for long drought periods, selection against the alternative allele would be desirable since the plant will keep stomata open, reducing the impact on photosynthesis activity and growth. Nevertheless, this selection must be done considering that a limited NTR would be necessary to avoid excessive loss of water. In this case, a positive selection for the alternative allele (A) of *Pv01*NTR_1 and a selection against the alternative allele (G) in *Pv01*NTR_2 should favor a reduced NRT.

The candidate gene *Phvul.001G108800* encodes an aquaporin SIP2-1-Related protein (Table 2). Aquaporins are small transmembrane channel proteins that facilitate the movement of water and small neutral molecules through pores across cell membranes (Maurel et al. 2015). In *Ath*, the aquaporin SIP2-1 protein is localized in the endoplasmic reticulum (ER) and has been associated with the alleviation of ER stress (Sato and Maeshima, 2019). ER stress results from the accumulation of misfolded or unfolded proteins in the ER during environmental stresses, including drought (Deng et al., 2013). Oocyte water transport essays conducted with the native sequence of the gene from common bean reference genome (PvSIP2) demonstrated its water channel activity (Figure 3). Sequence variation in the sequence of the gene may be advantageous in the hydraulic movement and water status of the plant in response to drought.

Thioredoxins (Trx) are thiol:disulfide oxidoreductase proteins involved in the formation or reduction of disulfide bonds (Eklund et al., 1991). This class of proteins can be induced by water deficit in *Solanum tuberosum* (Rey et al., 1998) and have been proposed as candidate genes for salt tolerance in bread wheat (Chaurasia et al., 2020). The candidate gene encoding for a Thioredoxin-Like 1-1 protein (*Phvul.001G109200*) may provide drought tolerance in common bean by regulating the redox state of chloroplast proteins. Recently, Yokochi et al. (2021) demonstrated in *Ath* that Trx and Trx-like proteins are involved in the regulation of chloroplast functions, including photosynthesis-related proteins.

*Pv01*NTR_2 had three candidate genes related to abiotic stress (Table 2). Extensins (EXTs) are proline-rich proteins that play a paramount role in cell wall synthesis and modification during root and pollen development (Kishor et al., 2015). Proline is an amino acid that accumulates under stress conditions as a beneficial solute to reduce the osmotic potential and help cope with the effects of drought (Chul et al., 2018). The candidate Phosphatidylinositol N-acetylglucosaminyltransferase subunit Q protein (PIG-Q) (*Phvul.001G127800*) is part of the GPI-GlcNAc-transferase (GPI-GnT) multiprotein complex involved in the glycosylphosphatidylinositol (GPI)-anchor biosynthesis in plants (Beihammer et al., 2020). The GPI-anchor is a post-transcriptional modification of diverse protein families implicated in cell wall composition, cell expansion, biotic and abiotic stress response, stomata development, pollen tube elongation, and hormone signaling (Beihammer et al., 2020). Variations in PIG-Q may change the modification of key proteins in the water deficit response. However, further investigation would be necessary.

Water stress increases reactive oxygen species (ROS) leading to oxidative stress. Nitric oxide (NO) limits the accumulation of ROS by inhibiting the Nicotinamide adenine dinucleotide phosphate (NADPH) oxidase enzyme complex, reacting with ROS, and inducing the expression of antioxidant enzymes genes (del Río, 2015). Furthermore, NO plays a key role in the abscisic acid (ABA)-induced stomatal closure during stress (Perlikowski et al., 2022). The nitric-oxide synthase (NOS) identified (candidate gene *Phvul.001G128000*) is one of the main enzymes involved in NO production (Neill et al., 2003) that may alleviate the oxidative stress and regulate an early stomatal closure. Cao et al. (2019) showed that an early NOS-mediated nitric oxidative increase mitigates oxidative damage in rice roots under drought.

Alkaline ceramidases have been poorly studied in plants. However, the overexpression of an alkaline ceramidase in *Ath* promotes autophagy under abiotic stresses (Zheng et al., 2018). Autophagy is a conserved pathway that plays a critical role in the adaptation of plants to drastically changing environmental stresses (Han et al., 2011). Particularly, the alkaline ceramidase AtACER is involved in disease resistance and salt tolerance (Wu et al., 2015). The Alkaline ceramidase-related candidate gene (*Phvul.007G090000*) is a novel target in common bean to study the role of alkaline ceramidases in crop plants, and also for drought tolerance improvement.

The transport of auxins plays an important role in plant growth and development, and is regulated by influx (AUXIN1/LIKE-AUX1 family) and efflux (mainly PIN-Formed (PIN)) carries (Swarup and Bhosale, 2019). The candidate Auxin efflux carrier component 1 (*Phvul.001G121100,* Table 2) is encoded by the PIN1 gene belonging to the type-I PIN proteins (Armengot et al., 2016). The down expression of PIN1 and other members of the type-I PIN subfamily has shown to affect root meristem size during salt stress by increasing NO levels (Korver et al., 2018). The potential role of PIN1 in common bean is in accordance with the candidate NOS gene detected also on Chr01. Furthermore, Zhang et al., (2012) showed that in rice PIN3t, another type-I PIN, is involved in drought stress response and tolerance.

On Pv07, the candidate gene *Phvul.007G088800* encodes the ATP-binding Cassette Transporter G family member 25 (ABCG25). ABCG25 is an ABA exporter expressed in vascular tissue that controls cellular ABA levels (Park et al., 2016) and influences stomatal regulation (Kuromori et al., 2010). ABA is the main plant hormone involved in abiotic stress responses, seed processes, and senescence. Particularly, long-distance ABA signaling and its interplay with other processes are involved in the detection of drying soil and plant responses (Davies et al., 2005), which agrees with our experiments. ABCG25 and other transporters have been shown to respond to abiotic stresses including temperature (Baron et al., 2012) and drought (Chen et al., 2011).

Phosphorylation is one of the main signaling mechanisms involved in a myriad of plant responses. The candidate genes *Phvul.007G089200* and *Phvul.007G089301* encoding for a Protein Tyrosine kinase (PTK) and a Calcium-dependent protein kinase (CPK) 11-related protein, respectively, may be involved in the response to drought stress. The role of PTKs has been mostly studied in animals and humans and remains poorly understood in plants. Recently, (Allimuthu et al., 2020) characterized a set of PTKs in rice, identifying key genes implicated in the osmotic and drought stress response. CPKs also phosphorylate Tyr residues of beta-tubulin and several transcription factors (TF) in *Ath* (Nemoto et al., 2015). Interestingly, many of these TFs are related to ethylene and ABA, and have shown to be relevant for plant response to abiotic stresses (reviewed in Miyamoto et al., 2019).

Five candidate genes involved in gene expression regulation were identified. Mitochondrial transcription termination factor (mTERF) proteins (*Phvul.007G089900*) are key components of organellar gene expression (OGE) machinery in chloroplasts and mitochondria (Barkan, 2011). Despite plants having more mTERF genes compared with animals, their role is still poorly understood. Studies in maize and *Ath* have shown that mTERFs regulate different developmental processes, including the response to abiotic stresses. For instance, in *Ath*, *mTERF* mutants display varying sensitivity to heat and saline stress, as well as response to ABA (Robles and Quesada, 2021). This poses an interesting role for this candidate gene in the drought tolerance of common bean.

The TFs bHLH106-related (*Phvul.001G121200*), C2H2-type Zn-finger protein (*Phvul.007G090500*), UPBEAT1 (*Phvul.007G089100*), and No apical meristem (NAM) protein (*Phvul.007G089600*) are promising candidate genes for drought tolerance improvement in common bean. bHLH TFs are one of the main TF families in angiosperms with paramount roles in abiotic stresses (Khan et al., 2018). In *Ath*, Ahmad et al. (2015) showed that bHLH106 encoded by the homologous gene *AT2G41130*, is a major regulator that integrates the function of multiple salt tolerance-related genes. However, in our validation experiments on *Ath*, *AT2G41130* was not involved in the regulation of the transpiration rate in response to drying soil (Figure 4).

C2H2 zinc finger proteins have key regulatory roles in abiotic stress resistance in plants. The CH2H-type proteins have a significant binding diversity characterized by their ability to interact with DNA, RNA, and other proteins (reviewed in Han et al., 2020). We showed that in *Ath*, the disruption of the homologous gene of *Phvul.007G090500* (*AT2G47270*) reduced the FTSWc and increased the transpiration rate post-FTSWc compared to the WT. This suggests the involvement of this gene in the regulation of stomata closure, delaying the moment and the fraction of water (FTSW) at which the plant closes the stomata. Natural allelic variation of this gene in common bean could be beneficial in response to water deficit conditions.

UPBEAT1 is a TF also belonging to the bHLH family. In *Ath*, UPBEAT1 regulates peroxidases and is involved in the accumulation of ROS and primary root elongation (Tsukagoshi et al., 2010). The mutant line *AT2G47270* showed consistent responses to drying soil. The knockout of the gene increased the FTSWc and NTR post-FTWSc compared to the WT suggesting a more conservative mechanism of water retention (Figure 4E). An earlier stomata closure was also accompanied by a slower reduction in the NTR after the FTSWc (lower slope; Figure 4F). While effects on plant growth may be observed with this type of strategy, in the event of long-term drought periods, this may be advantageous for plant survival.

Finally, the NAM proteins are part of the NAC TF family, which has been described in multiple crops (Shao et al., 2015), and associated with stress response in solanaceous (Tweneboah and Oh, 2017) and cereals (Dudhate et al., 2021).

## Conclusions

The present study represents the largest FTSW study reported. The high variation on mean NTR and FTSWc among Mesoamerican beans shows that variation for the improvement of these traits exists and may enable breeding cultivars with an improved capacity to preserve water status during short or long water deficit periods. The identified genotypes with the extreme values for mean NTR and FTSWc such as Othello, Harold, and ABCP-8 can be used as parents for future breeding strategies in either short or long drought periods. The four candidate QTL identified on Pv01 and Pv07 allowed the identification of 15 new candidate genes that can be targets for marker-assisted selection and for further research aimed at identifying causal polymorphisms in gene sequences. Interestingly, five candidate genes corresponded to transcription factors (e.g., *Phvul.007G089100*) which constitute major target genes due to their role in controlling multiple genes. We demonstrated the role of three candidate genes, one aquaporin and two transcription factors, in the transpiration rates of the plant in response to drying soil. Our results open the opportunity to exploit specific genes in common bean for the improvement of new drought-tolerant lines. Additionally, some of the candidate genes have been poorly studied in crop plans, leading to new insights into their potential physiological role in beans and other crop species.

## Notes

### Competing Interest Statement

The authors have declared no competing interest.

### Summary of Updates

Response: Thank you for your comment and insight. We hope that our additional research is sufficient to confirm the responses we have observed in bean. Although It is not a definitive account on the mechanisms, we feel that our work belongs in Plant Physiology as it spans various areas relevant to your readership, such as physiological mechanisms, genetics and breeding, molecular tools and bioinformatics. We feel it is also relevant to your readership as the results from our study have direct implications for the development of follow up field-based research approaches for the development of drought tolerance. Transpiration responses of plants to drying soil are key factors in studying the physiological basis of drought tolerance. We conducted a large-scale experiment analyzing the transpiration responses of common bean to drying soil. Candidate genes involved in the regulation of transpiration rates were identified through Genome-Wide Association (GWA) analyses, followed by functional studies to validate several of these candidate genes. Firstly, we examined the potential water channel activity of an aquaporin-related gene using oocyte swelling assays, synthesizing and expressing the native sequence of common bean. We demonstrated that the gene is involved in the water movement across membranes. Secondly, due to challenges in transformation and regeneration in common bean, we used mutant Arabidopsis plants to study homologous genes corresponding to the candidate genes. Using three Arabidopsis lines, we conducted two independent experiments replicating the conditions and analyses of transpiration rate measurements performed for common bean. Our results indicate that two out of the three evaluated transcription factors (TF) play roles in controlling transpiration in Arabidopsis. These findings provide novel insights into the physiological functions of the analyzed genes, with potential applications in crop improvement as well as in advancing understanding of the genetic control of water use in plants.

