## Supplementary materials for "Whole Plant Transpiration Responses of Common Bean (*Phaseolus vulgaris* L.) to Drying Soil: Water Channels and Transcription Factors"

**Table S1.** Primers used for the selection and corroboration T-DNA insertion in Arabidopsis mutant lines from ABRC for transpiration experiments.

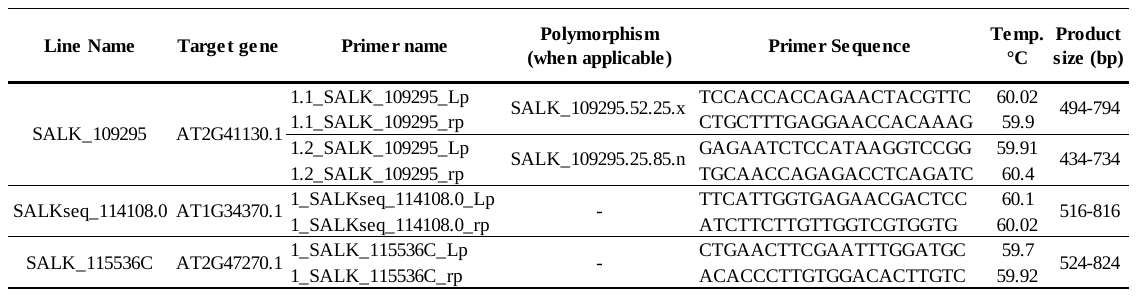

**Table S2.** Average Normalized Transpiration Rate (NTR) and critical Fraction of Transpirable Soil Water (FTSWc) the whole panel (DDP) of evaluated common bean genotypes. The R2 indicates the linear-plateau regression fitting to the data.

| **Genotype** | **Market Class** | **NTR** | **FTSWc** | | **R^2^** |
| --- | --- | --- | --- | --- | --- |
| Harold | Pink | 0.60 | 0.23 | a | 0.58 |
| CommonRedMexican | Small red | 0.54 | 0.26 | ab | 0.50 |
| ABCP_8 | Pinto | 0.96 | 0.26 | ab | 0.67 |
| Montrose | Pinto | 0.44 | 0.26 | ab | 0.83 |
| Buster | Pinto | 0.65 | 0.27 | ab | 0.37 |
| Poncho | Pinto | 0.48 | 0.27 | ab | 0.35 |
| Topaz | Pinto | 0.44 | 0.27 | ab | 0.65 |
| A285 | Cream | 0.71 | 0.30 | ab | 0.48 |
| Othello | Pinto | 0.37 | 0.30 | ab | 0.83 |
| Croissant | Pinto | 0.45 | 0.32 | ab | 0.57 |
| ShinyCrow | Black | 0.51 | 0.32 | ab | 0.56 |
| Matterhorn | Great Northern | 0.54 | 0.33 | ab | 0.55 |
| Stampede | Pinto | 0.53 | 0.34 | ab | 0.50 |
| ROG312 | Pink | 0.42 | 0.35 | ab | 0.73 |
| Quincy | Pinto | 0.48 | 0.35 | ab | 0.56 |
| IBC 301_204 | Small red | 0.56 | 0.36 | ab | 0.29 |
| Coyne | Great Northern | 0.56 | 0.36 | ab | 0.55 |
| BAT477 | Cream | 0.49 | 0.36 | ab | 0.50 |
| Buckskin | Pinto | 0.40 | 0.36 | ab | 0.59 |
| Marquis | Great Northern | 0.50 | 0.37 | ab | 0.62 |
| Weihing | Great Northern | 0.48 | 0.37 | ab | 0.48 |
| Fisher | Pinto | 0.42 | 0.37 | ab | 0.43 |
| UI_425 | Great Northern | 0.45 | 0.38 | ab | 0.33 |
| Avalanche | Navy | 0.53 | 0.39 | ab | 0.40 |
| MedicineHat | Pinto | 0.65 | 0.39 | ab | 0.36 |
| I9365_31 | Black | 0.52 | 0.39 | ab | 0.27 |
| Orion | Great Northern | 0.40 | 0.39 | ab | 0.76 |
| Yolano | Pink | 0.33 | 0.39 | ab | 0.65 |
| Kimberly | Pinto | 0.54 | 0.39 | ab | 0.36 |
| Verano | Small white | 0.51 | 0.39 | ab | 0.48 |
| Seafarer | Navy | 0.46 | 0.40 | ab | 0.47 |
| Mayflower | Navy | 0.55 | 0.40 | ab | 0.44 |
| NE2_09_3 | Pinto | 0.44 | 0.41 | ab | 0.52 |
| USPT_CBB_5_1 | Pinto | 0.59 | 0.42 | ab | 0.28 |
| USPT_CBB_1 | Pinto | 0.53 | 0.42 | ab | 0.41 |
| UI_59 | Great Northern | 0.38 | 0.43 | ab | 0.64 |
| BerylR | Great Northern | 0.57 | 0.43 | ab | 0.33 |
| US_1140 | Great Northern | 0.39 | 0.43 | ab | 0.46 |
| UI_537 | Pink | 0.51 | 0.43 | ab | 0.38 |
| PR0340_3_3_1 | Small red | 0.55 | 0.44 | ab | 0.41 |
| PR0443_151 | Black | 0.45 | 0.44 | ab | 0.35 |
| CDCCrocus | Great Northern | 0.48 | 0.44 | ab | 0.27 |
| Midnight | Black | 0.51 | 0.45 | ab | 0.37 |
| UI_239 | Small red | 0.66 | 0.46 | ab | 0.75 |
| Viva | Pink | 0.43 | 0.46 | ab | 0.57 |
| Zorro | Black | 0.44 | 0.46 | ab | 0.65 |
| LaPaz | Pinto | 0.50 | 0.46 | ab | 0.42 |
| Lariat | Pinto | 0.58 | 0.47 | ab | 0.35 |
| Sedona | Pink | 0.73 | 0.47 | ab | 0.34 |
| Chase | Pinto | 0.40 | 0.47 | ab | 0.56 |
| GN9_1 | Great Northern | 0.54 | 0.47 | ab | 0.39 |
| USRM_20 | Small red | 0.65 | 0.47 | ab | 0.32 |
| Schooner | Navy | 0.55 | 0.47 | ab | 0.45 |
| SEA10 | Cream | 0.63 | 0.47 | ab | 0.60 |
| GN9_4 | Great Northern | 0.51 | 0.48 | ab | 0.39 |
| C_20 | Navy | 0.47 | 0.48 | ab | 0.46 |
| Victor | Pink | 0.73 | 0.48 | ab | 0.52 |
| Common Pinto | Pinto | 0.64 | 0.49 | ab | 0.47 |
| PT7_2 | Pinto | 0.47 | 0.50 | ab | 0.27 |
| 115M (BlackRhino) | Black | 0.70 | 0.50 | ab | 0.38 |
| A_55 | Black | 0.82 | 0.51 | ab | 0.29 |
| Roza | Pink | 0.48 | 0.51 | ab | 0.43 |
| PinkFloyd | Pink | 0.57 | 0.52 | ab | 0.37 |
| BillZ | Pinto | 0.40 | 0.52 | ab | 0.60 |
| Raven | Black | 0.54 | 0.52 | ab | 0.55 |
| Medalist | Navy | 0.41 | 0.53 | ab | 0.56 |
| BelNeb_RR_1 | Great Northern | 0.56 | 0.53 | ab | 0.41 |
| Gemini | Great Northern | 0.42 | 0.54 | ab | 0.71 |
| PT9_17 | Pinto | 0.43 | 0.54 | ab | 0.52 |
| TARS_VCI_4B | Pinto | 0.38 | 0.55 | ab | 0.52 |
| Maverick | Pinto | 0.47 | 0.56 | ab | 0.39 |
| F04_2801_4_1_2 | Black | 0.46 | 0.57 | ab | 0.55 |
| Merlot | Red | 0.57 | 0.58 | ab | 0.49 |
| SantaFe | Pinto | 0.52 | 0.59 | ab | 0.42 |
| Gloria | Pink | 0.61 | 0.59 | ab | 0.61 |
| Kodiak | Pinto | 0.51 | 0.60 | ab | 0.27 |
| DOR 364 | Small red | 0.65 | 0.61 | ab | 0.45 |
| Nodak | Pinto | 0.45 | 0.62 | ab | 0.29 |
| CENTAPupil | Small red | 0.43 | 0.63 | ab | 0.53 |
| F07_449_9_3 | Red | 0.42 | 0.66 | ab | 0.33 |
| NW_63 | Small red | 0.43 | 0.64 | ab | 0.37 |
| T_39 | Black | 0.43 | 0.99 | b | 0.70 |
| **Mean** | | 0.52 | 0.45 | | 0.48 |

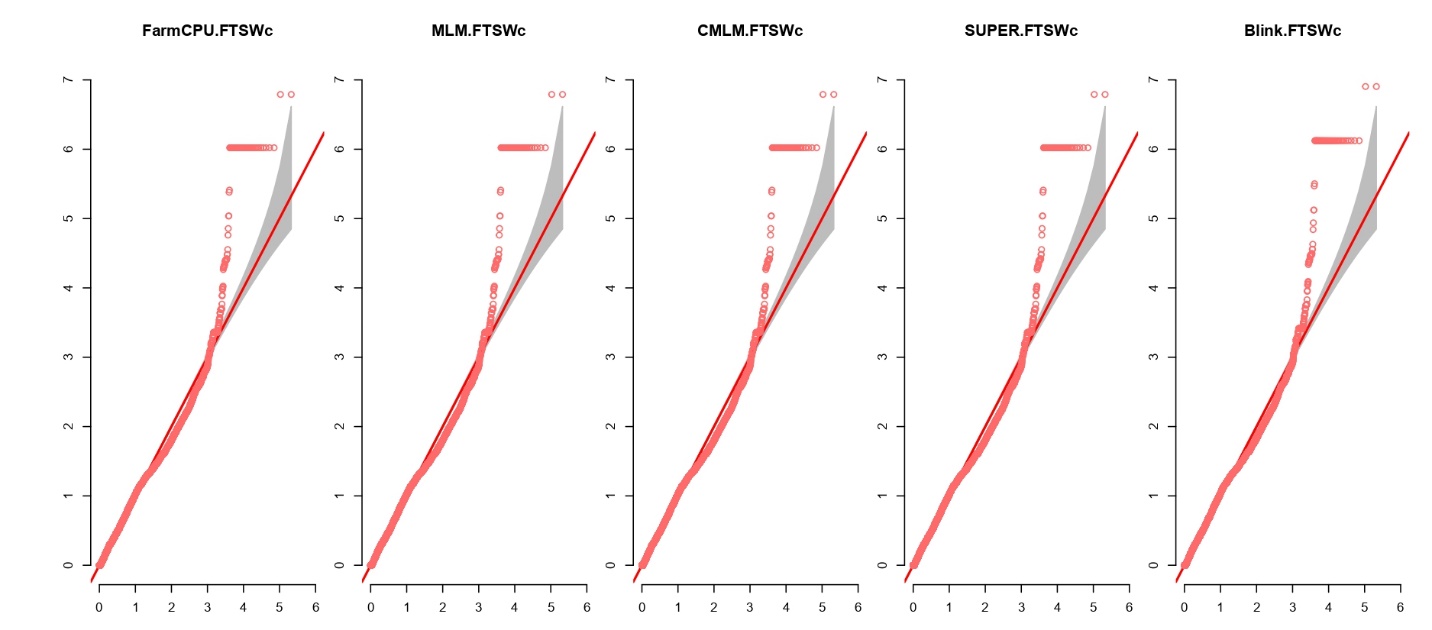

**Figure S1.** Fitting of different GWAS models for critical Fraction of Transpirable Soil Water (FTSWc). Fixed and random model Circulating Probability Unification (FarmCPU), Mixed Linear Model (MLM), compressed MLM (CMLM), Settlement of MLM Under Progressively Exclusive Relationshipc(SUPER), Bayesian-information and Linkage-disequilibrium Iteratively Nested Keyway (Blink).

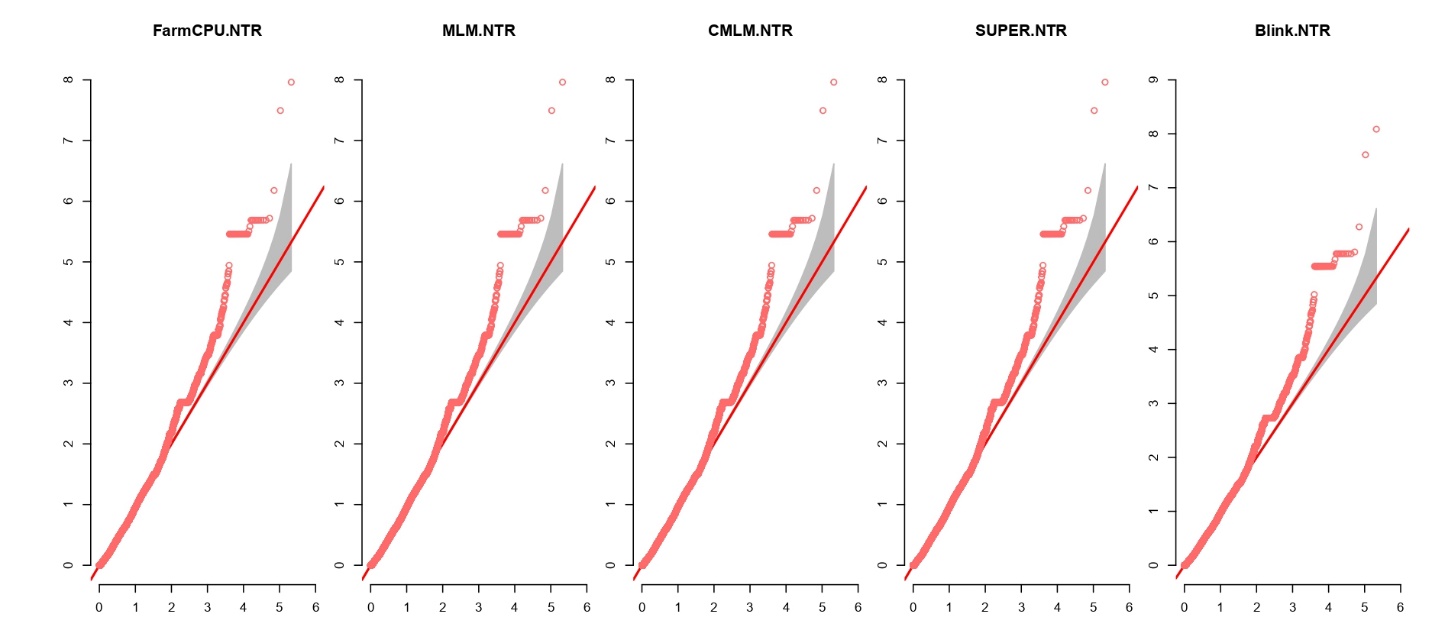

**Figure S2.** Fitting of different GWAS models for critical Average Normalized Transpiration Rate (NTR). Model descriptions as above.
